# Origin of pest lineages of the Colorado potato beetle, *Leptinotarsa decemlineata*

**DOI:** 10.1101/156612

**Authors:** Victor Izzo, Yolanda H. Chen, Sean D. Schoville, Cong Wang, David J. Hawthorne

## Abstract

Colorado potato beetle (*Leptinotarsa decemlineata* Say) is a pest of potato throughout the Northern Hemisphere, but little is known about the beetle’s origins as a pest. To determine the origins of pest populations of CPB, we sampled the beetle from uncultivated *Solanum* host plants in Mexico, and from pest and non-pest populations in the U.S. We used mtDNA and nuclear loci to examine three hypotheses on the origin of the pest lineages: 1) the pest beetles originated from Mexican populations, 2) the pest beetles descended from hybridization between previously divergent populations, or 3) the pest beetles descended from populations that are native to the Plains states in the United States. We examined patterns of genetic diversity among geographic regions in order to detect invasion-related genetic information. Mitochondrial haplotypes of non-pest populations from Mexico and southern Arizona differed substantially from beetles collected from the southern plains and potato fields in the U. S., indicating that beetles from Mexico and Arizona did not contribute to founding the pest lineages. Similar results were observed for AFLP and microsatellite data. In contrast, non-pest populations from the southern plains of Colorado, Kansas, Nebraska, New Mexico, and Texas were genetically similar to U. S. pest populations, clearly indicating that they contributed to the founding of the pest lineages. Although some pest populations are less genetically diverse (e.g., Washington, Idaho), most of the pest populations do not show a significant reduction in genetic diversity compared to the plains populations in the U. S. In contrast to the colonization patterns typical of exotic pests, our analyses suggests that a large genetically heterogeneous beetle populations expanded onto potato from native *Solanum* hosts. As an endemic colonization of a novel host plant, this host range expansion may have contributed to the relatively abundant genetic diversity of contemporary populations, perhaps contributing to the rapid evolution of host range and insecticide resistance in this widely successful insect pest.

## INTRODUCTION

The Colorado potato beetle (CPB), *Leptinotarsa decemlineata* Say (Coleoptera: Chrysomelidae) is one of the most damaging insect pests of potato (Weber 2003). The beetle displays a remarkable ability to adapt to abiotic and biotic stressors, making it an excellent model for the study of rapid evolution. CPB has adapted to: a wide range of host plants within the plant family Solanaceae (Hsiao 1985, Hare and Kennedy 1986), over 50 different insecticides in all the major classes (Casagrande 1987, Alyokhin et al. 2008) and diverse climatological and phenological regimes (Tauber et al. 1988, Lehmann 2013, Izzo et al. 2014, Lehmann et al. 2015). Although many studies describe the rapid evolutionary dynamics of this beetle the origins of the pest lineages and relationships among beetle populations remain obscure (Hsiao 1985, Hare and Kennedy 1986, Hare 1990, Lu et al. 2001, Piiroinen et al. 2013, Izzo et al. 2014). A species’ ability to respond to environmental variation hinges on the distribution of genotypic and phenotypic variation within and among populations, which may be significantly impacted by its phylogeographic histories of those populations (Slatkin 1987, Roderick 1996, Chen et al. 2006). Boiteau (1994) suggested that CPB exhibits moderately high levels of genetic variation at nuclear loci across the North American portion of its invaded range, an observation supported by subsequent population genetic analyses (Hawthorne 2001, Grapputo et al. 2005, Piiroinen et al. 2013). Understanding current patterns of population genetic variation and the relationships among geographic CPB populations may reveal their roles in colonization of temperate potato agroecosystems and contribute to our understanding of how CPB emerged as a successful global pest.

CPB is endemic in North America, where it originally fed on several plant species in the genus *Solanum*, namely *S. angustifolium*, *S. rostratum*, and *S. eleagnifolium* (Tower 1906, Hsiao 1981, Jacques 1988). First collected in the U.S. in 1811 along the Nebraska and Iowa border, most likely on *S. rostratum* (Casagrande 1985), CPB was not recognized as a pest until it expanded its host range to include potato, *S. tuberosum* L. in 1859 (Walsh 1865). Subsequently, CPB became a devastating insect pest of potatoes, spreading from the Central Plains to the East Coast of the U.S. in only 15 years, and to the Western states and provinces of the US and Canada by the 1920s (Tower 1906, Gauthier et al. 1981, Hsiao 1985). Despite the detailed historical record of the CPB’s geographic expansion across the North American continent (Walsh 1865, Tower 1906, Jacques 1988), the origin of pest CPB populations remains unknown. Here, we pose and evaluate three hypotheses of distinct colonization pathways of pest populations of CPB based on patterns of genetic variation.

The majority of *Leptinotarsa* species (including *L. decemlineata*) are found within Mexico (Jacques 1988) and initial records of CPB are from the U.S. (Tower 1906, Casagrande 1987, Hare 1990). These observations suggest that central Mexico is the most likely ancestral region for the species, and the recent origin of pest populations (Tower 1906, Hsiao 1978, Logan et al. 1987, Jacques 1988, Hare 1990). The “Out-of-Mexico” hypothesis suggests that 16^th^ century Spanish expeditions and subsequent cattle ranching, inadvertently expanded the range of the beetle’s host plant, *S. rostratum* during their northern migration into the current western U.S. (Hsiao 1981, Jacobson and Hsiao 1983, Casagrande 1985, 1987, Lu and Lazell 1996). The northward expansion of *S. rostratum* as a rangeland weed then facilitated the subsequent latitudinal expansion of the CPB into the U.S. (Tower 1906, Lu and Lazell 1996, Zhao et al. 2013). Because the beetle is not a pest of potatoes within Mexico, we suggest that the potato feeding phenotype of the beetle did not emerge until it encountered potato in the central U.S. (J. Núñez-Farfán, pers. comm., V. Izzo, pers. obs., Y. H. Chen, pers. obs). The “Out-of-Mexico” hypothesis would result in distinctive patterns of CPB genetic diversity in N. America (Table 1). Under the “Out-of-Mexico” hypothesis, we would expect that CPB in the U.S. (both Pest and non-pest samples from *S. rostratum* and *S. eleagnifolium*, “Plains” samples) would have mitochondrial and nuclear haplotypes that were similar to those found in Central or Northern Mexico, as they have been separated for only approximately 400 years. Also, the sampling effects of genetic drift that occurred during the range expansion would lead to reduced genetic diversity in the Plains and Pest samples relative to Mexican samples (Sakai et al. 2001).

**Table 1.**
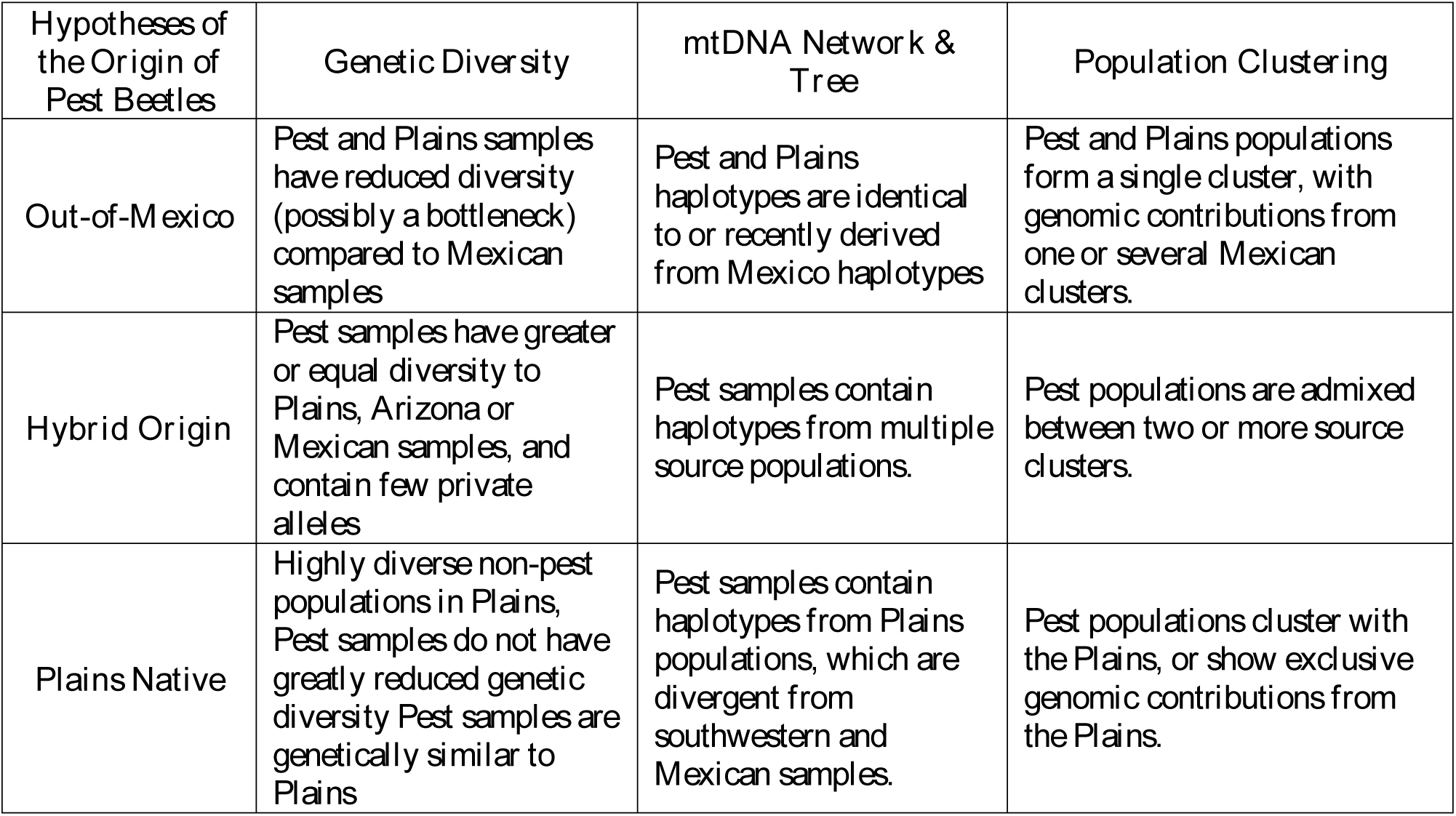
Three hypotheses for the origin of genetically variable pest populations of Colorado potato beetle and the genetic evidence that would support each.

Hsiao (1985) first suggested the “Hybrid Origin” hypothesis following analysis of chromosomal inversions found to be associated with pest geography. He hypothesized that pest populations are genetically hyper-diverse because they were founded via a recent hybridization between geographically separated and genealogically divergent races of CPB (Hsiao 1985). Using karyotyping, he found that desert CPB populations in northern Mexico, Arizona and Utah had a metacentric inversion on the second chromosome, while populations in southern Texas had an acrocentric inversion. Surprisingly, he also discovered that many U.S. pest populations carried both inversion types. Hsiao suggested that southwestern U.S. populations had mixed with another source population to yield a hybrid pest population with inflated genetic diversity. If pest beetles are the result of admixture of previously geographically-isolated populations, including ‘ghost populations’ that are extinct or unsampled, contemporary CPB pest populations should show a mixture of relatively divergent genotypes at many loci. Under the Hybrid Origin hypothesis, genetic assays using nuclear loci would reveal greater genetic diversity in Pest samples compared to Plains samples. Furthermore, pest populations would be expected to be intermediate to the source types in genetic distance and in phylogenetic position (Roderick 2004). Under the Hybrid Origin hypothesis, pest populations would also be expected to have mitochondrial DNA (mtDNA) haplotypes from divergent sources (Table 1).

Walsh (1865) proposed the third hypothesis that CPB is native to the eastern slope of the Rockies, where it fed on *S. rostratum* long prior to the arrival of Europeans and the potato plant. Under this “Plains Native hypothesis”, endemic pest populations would retain considerable genetic variation as they colonized potato within their native range and they are expected to be very genetically similar to plains source populations. Support for the “Plains Native hypothesis” scenario would include: high levels of genetic diversity among non-pest samples of CPB in the central and southern plains of the U.S., which are also potentially divergent from Mexican samples reflecting their long residence in the plains, significant genetic similarity of non-pest Plains and Pest populations, and sizable genetic diversity in pest populations indicating the lack of a colonization-mediated genetic bottleneck (Table 1).

Previous molecular studies utilizing lower-resolution tools such as allozymes (Jacobson and Hsiao 1983) and restriction fragment analysis of mitochondrial DNA (Azerado-Espin et al. 1991, Zehnder et al. 1992) have provided limited information on the origin of the CPB pest populations and display conflicting results. Jacobson and Hsiao (1983) and Zehnder et al. (1992) revealed limited genetic variation in CPB north of Mexico, whereas Azerado-Espin et al. (1996) found considerable variation. Most notably, the limited sampling of CPB from Mexican and southwestern U.S. locations in the previous population genetic studies precludes testing the “Out-of-Mexico” and Hybrid origin hypotheses.

In this study, we tested hypotheses outlined in Table 1 on the founding of the pest populations of CPB using a geographically wider sampling scheme and higher resolution genetic tools than previous studies. We collected beetle populations across a large number of geographic locations, and we genotyped them using mtDNA and nuclear loci. Reconstruction of the ancestry of the pest genotypes will help clarify the evolutionary processes that occurred during CPB’s colonization of temperate potato agroecosystems and may provide insight on how this species continually evolves as it spreads to new environments and faces new insecticides.

## MATERIALS AND METHODS

### Sample collection and preparation

We collected samples of *L. decemlineata* during two separate periods over a course of 14 years. We labelled samples according to the states from which they were collected with the exception of a sample from Long Island, NY, which was labelled separately (LI) from those sampled from the remainder of New York. In most cases, we collected from multiple sampling areas in each state. We defined a sampling area as a stand of host plants located at least 5 km apart. Within sampling areas, we collected individual beetles from plants located at least 1 m apart in order to sample the most diverse collection possible. After collection, we placed insect specimens directly into 95% ethanol and stored them at −20⏢. Following collection, we froze individual beetles in liquid nitrogen and stored them at −80⏢. DNA was extracted using the DNeasy Tissue Kit (QIAGEN), following the protocol for animal tissues. We analyzed two sets of CPB specimens collected over two time-periods: 1998 and 2010-2012.

### Sampling Period 1 (1998)

We collected 140 beetles from across the range of *L. decemlineata* within the U.S. and refer to this sampling assemblage as SP1 (Figure 1, Table S.1) These samples represented pest beetles collected from commercial potato (*Solanum tuberosum*) and non-pest individuals collected from native *Solanum spp.* (*S. rostratum* or *S. eleagnifolium*). Pest beetles were collected from commercial potato fields in Great Lakes and Atlantic coast states and also in two northwestern locations of the U.S. Samples from non-pest populations were collected along roadways and near livestock pens in the Great Plains region of the United States (Colorado, Kansas, Nebraska, New Mexico and Texas — hereafter named “Plains” samples), and in southern Arizona. Although one Plains sample (Springlake, TX) was collected near a potato field, potato production is uncommon in the region where the Plains individuals were collected, and there is no commercial potato production in southern Arizona.

**Figure 1.**
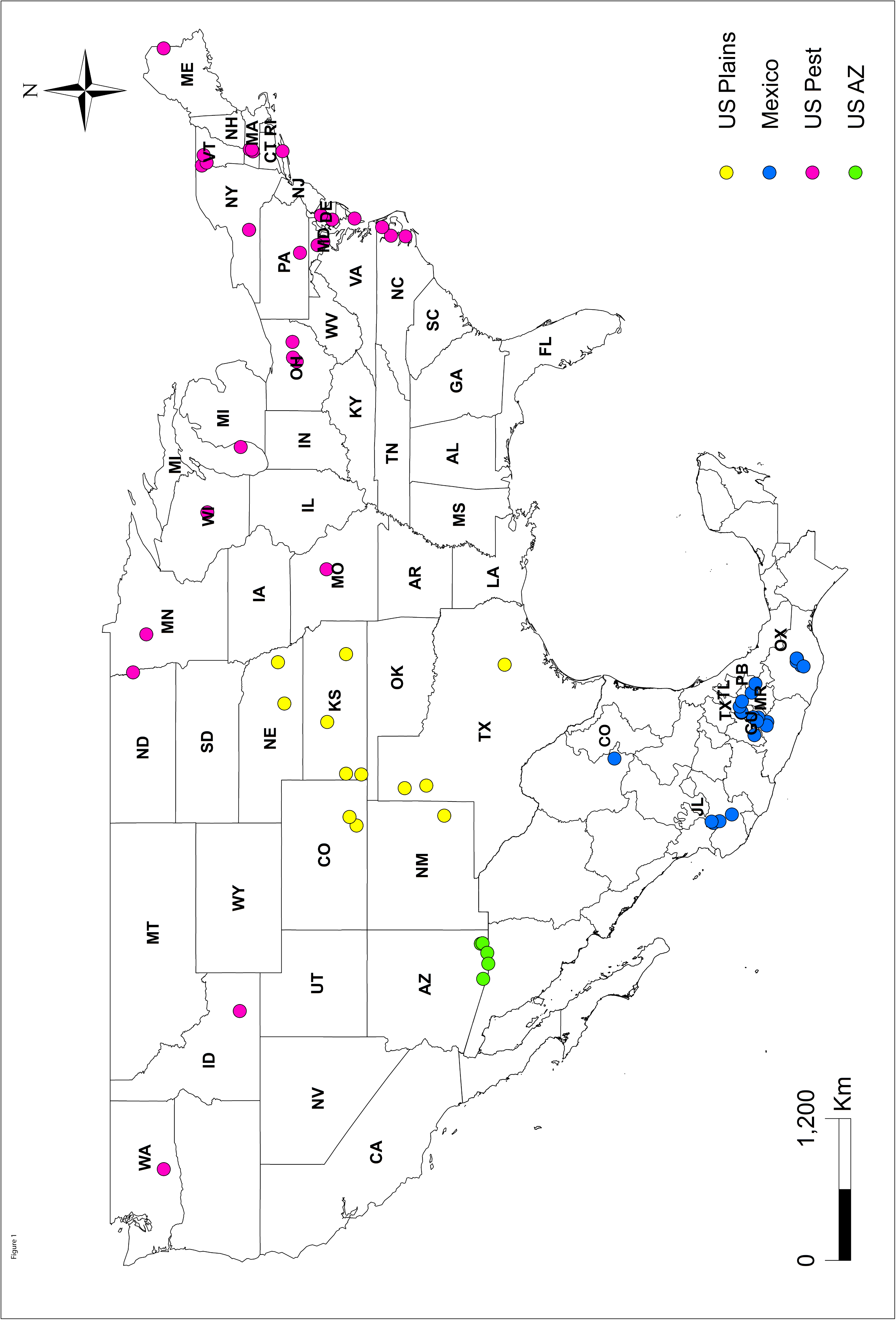
Sampling locations of Colorado potato beetle for SP1 and SP2.

### Sampling period 2 (2010-12)

In the second sampling period (SP2), we collected 400 *L. decemlineata* from locations distributed throughout the U.S. and Mexico (Figure 1, Table S. 1). Similar to SP1, we collected beetles from both pest and non-pest populations. All non-pest beetle samples collected in Mexico were collected from wild *Solanum* species (*S. rostratum* and *S. angustifolium*) typically found on roadsides throughout southern/central Mexico. Beetles were sampled from the Mexican states of Jalisco (JL), Morelos (MR), Puebla (PB), Texcoco (TX), Coahuila (CO), and Tlaxcala (TL) from *S. rostratum*. We sampled beetles from *S. angustifolium* in the Mexican states of Guerrero (GU) and Oaxaca (OX). Pest beetles in SP2 were sampled from within commercial potato growing fields at 6 locations in the U.S., including Vermont, Massachusetts, Maryland, Virginia, Washington and Missouri, and two non-pest plains locations sampled from *S. rostratum* in Texas and Kansas.

### mtDNA from SP1 and SP2

A 577-bp fragment of mtDNA spanning the 3′ end of the COI gene to the 5′ end of the COII gene using the primers S2792 (Brower 1994) and C2-N-3389 (Simon et al. 1994) was amplified from a subset of the SP1 samples and from all of the SP2 samples. An additional sample of *Leptinotarsa juncta* from College Park, MD was included in the mtDNA sequence analyses. Following amplification, fragments were commercially purified and sequenced in both forward and reverse reactions using fluorescently labeled primers (see above) on ABI 3730xl DNA analyzers by Genewiz (South Plainfield, NJ, SP1 samples) or by the High Throughput Genomics Center (University of Washington, Seattle, WA, SP2 samples).

We aligned 337 mtDNA sequences using ClustalW in Mega 6.0 (Tamura et al. 2013). We trimmed and edited sequences for quality resulting in fragments with a common length of 546 bp. Sequence polymorphism statistics were estimated for each sample location, including nucleotide diversity (π, the average number of nucleotide substitutions per site between sequences) and (H), haplotype diversity (Nei 1987), using DnaSP 5.10.1 (Librado and Rozas 2009).

We tested relationships among haplotypes as a median joining network with the default settings and the MP option using the PopART software (http://popart.otago.ac.nz, Bandelt et al. 1999, Polzin and Daneshmand 2003) Finally, to further illustrate phylogeographic relationships among sampled regions, we constructed a Bayesian phylogenetic tree as estimated in the program BEAST2 v2.4.0 (Bouckaert et al. 2014). Our priors included the HKY model with a gamma correction, as this was favored as a top performing model in the model test function of MEGA 6.0 (Tamura et al. 2013). Additionally, we selected a Yule model as a tree prior, a strict clock, and ran two independent runs for 100 million generations. Both runs were assessed for convergence and high ESS values (>200) in TRACER, then combined to estimate a maximum clade credibility tree. We used Figtree to report the consensus tree and posterior probability support values.

### AFLP (SP1)

We generated AFLP constructs of *Pst*1 and *Eco*R1 fragments from digested DNA and amplified them as described in Hawthorne (2001). Nine amplifications were performed (primer combinations in Table S.2) and silver stained fragments were visualized on a light box, recorded as present or absent and digitally archived by scanning the gel directly. We used a binary categories for scoring the gels (1 = band present, 0 = absent) and discarded markers with unresolved discrepancies. For each individual we arrayed the 0-1 score for each of the bands and analyzed them as a multilocus haplotype. Because a significant negative correlation of fragment frequency and fragment size may be caused by excessive homoplasy, we estimated that correlation using AFLP-SURV (VEKEMANS 2002).

We calculated the Euclidian distances among haplotypes using ARLEQUIN V3.0 (Excoffier et al. 2007) and used AMOVA to partition the AFLP genotypic variance into within and among sampling site components. Pairwise comparisons among sample sites yielded a matrix of average genetic distances among populations (Nei 1987). AFLP fragments were analyzed as recessive markers in STRUCTURE v2.3.4 (Falush et al. 2007) to identify population clusters and individual admixture proportions. We used a burn-in of length 50,000 and tested values of *K* = 2 – 7 groups. STRUCTURE runs were performed under an admixed model of ancestry and the correlated allele frequency model with 50,000 Monte Carlo Markov-Chain (MCMC) repetitions. Because samples from the Pacific Northwest (WA & ID) had limited AFLP variation and led to unusual geographical patterns of cluster membership, we examined the effect of removing these samples in a second STRUCTURE analysis. STRUCTURE runs were replicated ten times for each value of K, and the “best” K was inferred using the methods of Evanno *et al.* (2005) and figures created from the aggregated replicates using Clumpak (Kopelman et al. 2015).

AFLP markers were analyzed as individual loci using the Bayesian methods of Zhivotivsky (1999) to estimate allele frequencies and methods of Lynch and Milligan (1994) to estimate genotype frequencies in AFLP-surv. We compared the levels of genetic diversity across the Pest, Plains and Arizona groupings identified by structure analysis of AFLP. Nei’s genetic diversity (H) and genetic distances among samples were estimated using AFLP-surv.

### Microsatellites (SP2)

Using the SP2 samples, we genotyped 16-32 individuals in each of 8 sample locations in the US and 7 in Mexico, for a total of 400 sampled individuals, using six microsatellite loci reported by Grapputo et al. (2006). We followed the PCR protocols of Grapputo et al. (2006) to amplify the microsatellite loci with fluorescently labeled primers. PCR products were separated via capillary electrophoresis on an ABI Prism 3130xl Genetic Analyzer (Applied Biosystems, Foster City, CA, USA) at the University of Vermont Cancer Center. We manually binned microsatellite size classes using the software PeakScanner (Applied Biosystems) and constructed genotype profiles for advanced analysis. We used MICRO-CHECKER to correct allele frequencies that may have a deficit of heterozygotes due to PCR artifacts (Van Oosterhout et al. 2004). Genotype frequencies were adjusted for null allele artifacts in each of the SP2 sample locations before use in subsequent analysis.

We evaluated per-locus and per-population genetic diversity on the basis of allelic richness (RS, number of alleles independent of sample size), observed (H_O_) and expected (H_E_) heterozygosity, inbreeding coefficient (F_IS_), and linkage disequilibrium. To compare population-level relationships identified by microsatellite markers, we constructed neighbor-joining (NJ) trees based on pairwise genetic distances. For each population pair, we calculated Nei’s standard genetic distance (Nei 1987). All calculations were performed in FSTAT (Goudet 1995) with significance based on Bonferroni corrected *p* values after 10,000 random permutations.

As with AFLPs, we used STRUCTURE to assess clustering of individuals based on microsatellite genotypes, using a burn-in of length 100,000 and *K* = 1 – 15 groups. The STRUCTURE runs used an admixed model of ancestry and the correlated allele frequency model with 100,000 Monte Carlo Markov-Chain (MCMC) repetitions. The number of clusters, “K” was inferred from ten replicates of each “K” and figures created from the replicates using Distruct (Rosenberg *et al*. 2001).

## Results

### Mitochondrial Sequence Analysis

We identified 73 mtDNA haplotypes from 341 CPB individuals, combined from our SP1 and SP2 datasets (Figures 2 & 3). Of the 577 nucleotides sequenced from the COI-COII fragment, 76 (13.2%) were polymorphic. Both the Bayesian phylogeny (Figure 2) and the minimum spanning network (Figure 3) clearly show a large and discrete break between the northern (Pest + Plains) and the southern (Arizona + Mexico) samples, with 15 substitutions between the nearest Pest + Plains and Arizona + Mexico haplotypes. In both analyses, the portion of the tree or network including the Arizona and Mexico samples shows three coherent clusters of haplotypes from different geographic locations, with GU, JL, MR and OX in one, TX, TL, and PB in another and the more northern Arizona and Coahuila samples in the third.

**Figure 2.**
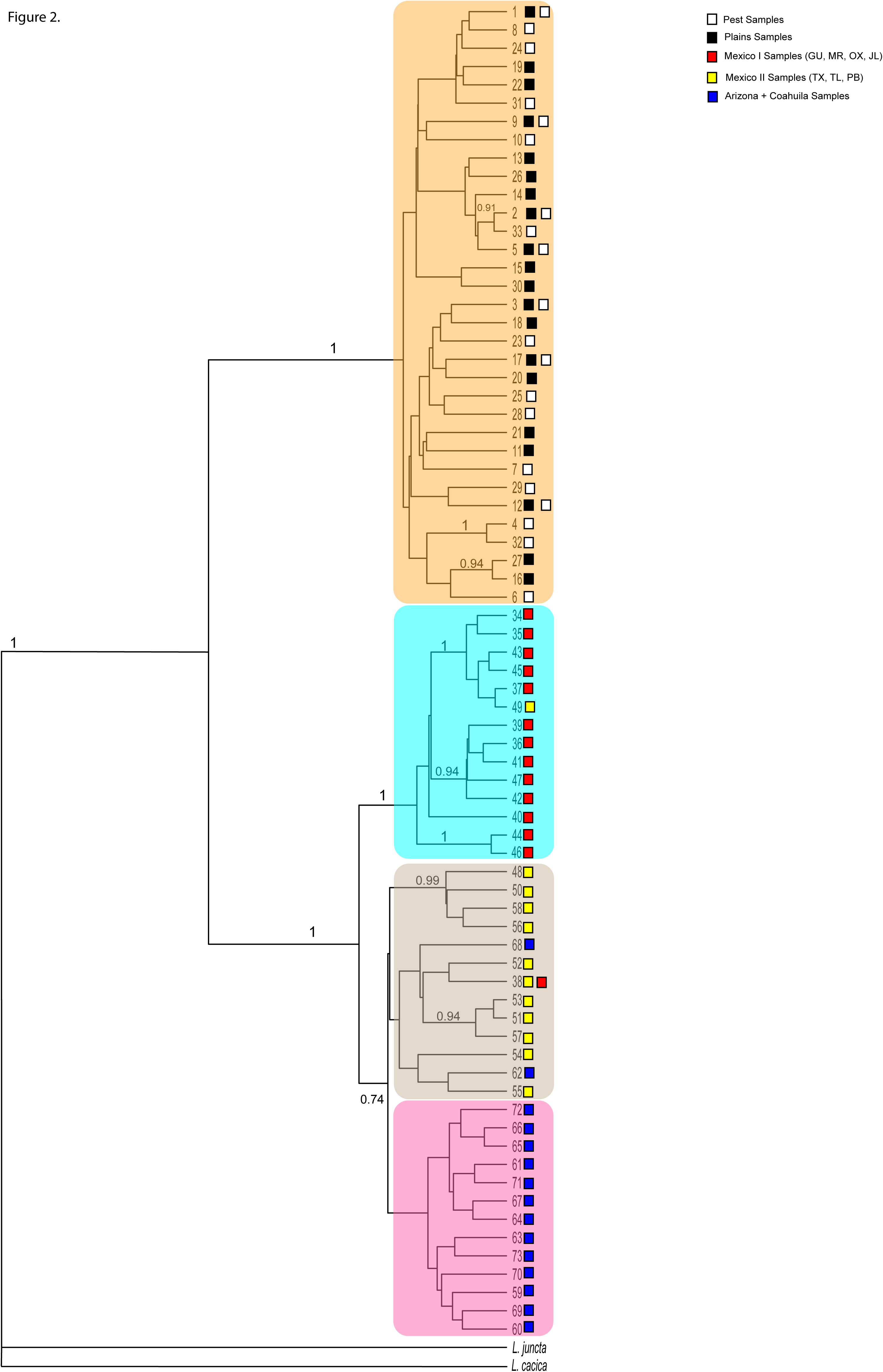
Phylogram of mtDNA haplotypes from Colorado potato beetle (*L. decemlineata*) and two related species (*L. juncta, L. cacica*). Posterior support values for clades are on branches, colors delineate geographic clades and boxes at the tips are colored to represent sample regions.

**Figure 3.**
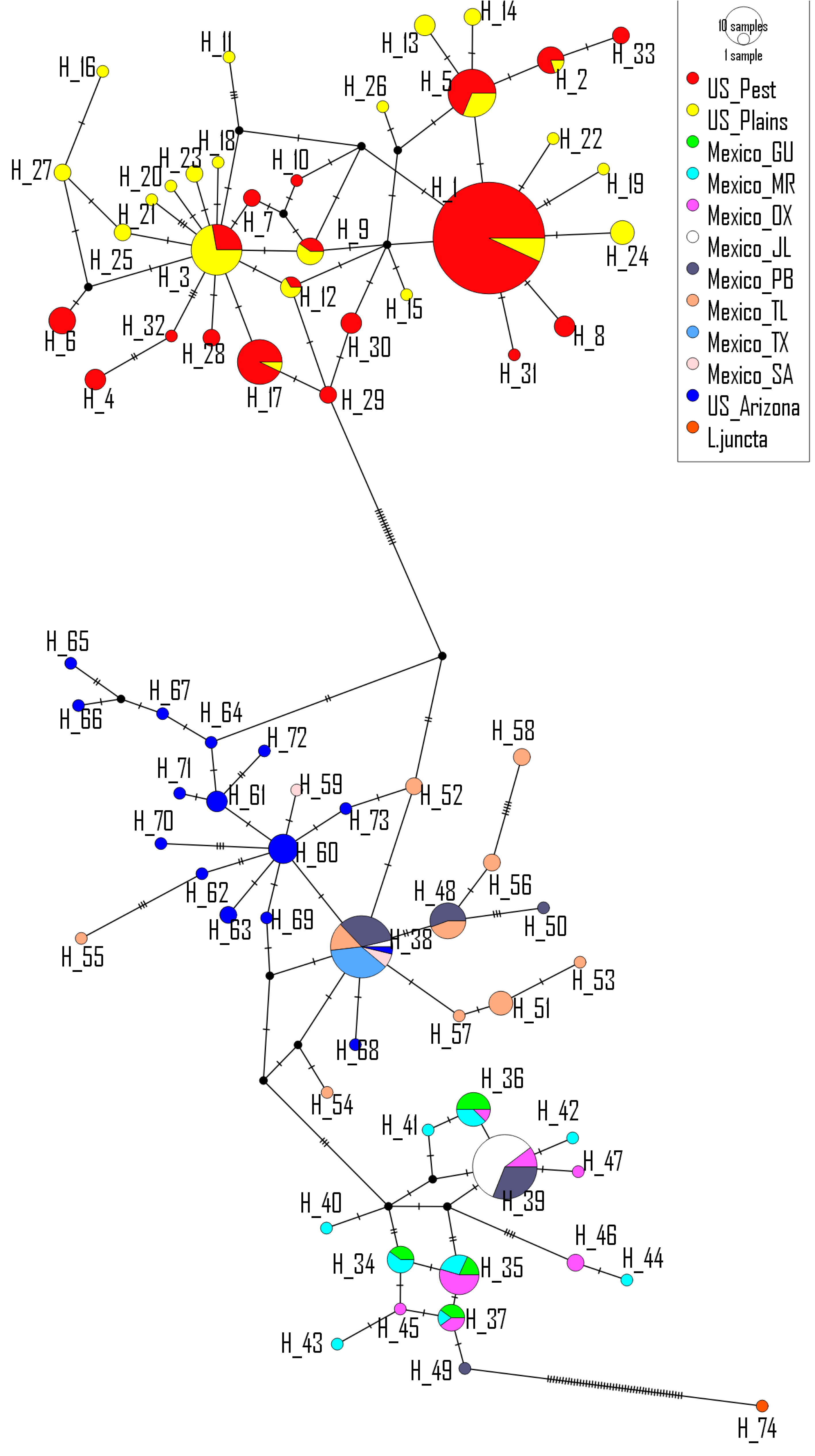
Network diagram of mtDNA haplotypes sampled from Colorado potato beetle. Hash marks represent nucleotide substitutions between nodes. Nodes are labeled with haplotype numbers. Unlabeled nodes are inferred but unsampled haplotypes.

The four most common pest haplotypes (1, 3, 5 and 17) were shared by Pest and Plains samples dispersed much more broadly (LI, MA, ME, MI, MN, MD, NE, NY, OH, PA, TX, WI) than any non-pest haplotypes and (Figure 2). Within the Arizona and Mexico sub-cluster, a single haplotype (38) was shared among the Arizona, Coahuila and the JL, TX, TL, and PB samples, and one haplotype (39) was shared among samples from the two southern Mexico clusters. We did not find clusters separating the Pest or Plains haplotypes. Haplotype 1 occurs in a large and geographically diverse collection of Pest and Plains samples, among which six additional haplotypes were clearly shared among Pest and Plains samples (Figure 3). Samples from the Pacific Northwestern areas of the U.S., Washington and Idaho, had reduced haplotype diversity, with all 11 WA and ID samples sharing haplotype 5, which was also shared with Pest beetles from Massachusetts and Plains beetles from Kansas and Texas (Figure 3).

### Diversity and Divergence measures

The complete set of samples showed substantial genetic diversity with a haplotype diversity (H) of 0.911 and nucleotide diversity (π) of 0.024 (Table 2). Seventeen haplotypes were found in the U.S. sample of 128 Pest beetles (10 were unique to pest samples), 24 haplotypes were found in the Plains sample (N = 69) (of which 15 were unique to plains samples), 15 haplotypes were found in the Arizona sample (N = 23) and 26 haplotypes were found among the samples from Mexico (N = 123). The combined Mexico + Arizona samples displayed greater overall haplotype and nucleotide diversity than the Pest or Plains samples with a haplotypic diversity (H) of 0.910, and an average nucleotide diversity (π) of 0.011. When Mexican sample locations are divided into the groups indicated by the mtDNA network analysis (Group 1 = GU, MR, OX, JL; Group 2 = TL, TX, PB), each of the two groups contained less haplotype diversity (H) (Group 1 = 0.780, Group 2 = 0.736), and nucleotide diversity (π) (Group 1 = 0.0050, Group 2 = 0.0050). Plains samples had slightly less haplotype diversity (H = 0.892) than the combined Mexican samples but more than either of the two individual Mexican groups. Nucleotide diversity of the Plains samples (π = 0.0052) was less than half of the combined Mexico + Arizona samples, but similar to individual Arizona (π = 0.0050) or Mexican groups. The Pest samples of CPB had slightly reduced haplotype and nucleotide diversities (H = 0.66; π = 0.0041), compared to the Plains, Arizona or Mexican sample groups (Table 2). AMOVA attributed 77.0% of the COI-COII haplotype variation to variation among the Pest-Plains-Mexico + Arizona groupings, 11.3% to variation among the state or regional level samples within those groupings and 11.8% to variation within those samples. Nucleotide divergence (uncorrected for intra-regional variation) between Pest and Mexico + Arizona haplotypes averaged 4.3% and between Plains and Mexico + Arizona the mean divergence was 4.2% (Table 3).

**Table 2.** mtDNA diversity indices from N. American CPB. Sample sets represent groupings used for hypothesis testing. “Whole sample” includes all successfully sequenced individuals sampled during the study. “Mex + AZ” includes all beetles sampled from Mexico and Arizona. “Plains” samples are from southern plains of U.S. collected from *S. rostratum*. “Mex” represents Mexican beetles. “Mex1_GUMROXJL” includes sequences from the Mexican states of Guerrero, Morelos, Oaxaca, and Jalisco. “Mex2_TLTXPB” includes sequences from the Mexican states of Tlaxcala, State of Mexico, and Puebla. “AZ” represents samples from Arizona.

| Sample Sets | N | #Polymorphic sites | #Haplotypes | Haplotype Diversity (h) | Nucleotide Diversity (pi) |
| --- | --- | --- | --- | --- | --- |
| Whole Sample | 337 | 76 | 72 | 0.911 | 0.0242 |
| Mex+AZ | 140 | 43 | 39 | 0.910 | 0.0110 |
| Plains | 69 | 24 | 23 | 0.892 | 0.0052 |
| Pest | 128 | 18 | 17 | 0.656 | 0.0041 |
| Mex | 117 | 32 | 25 | 0.877 | 0.0103 |
| Mex1_GUMROXJL | 69 | 19 | 14 | 0.780 | 0.0050 |
| Mex2_TLTXPB | 48 | 27 | 12 | 0.736 | 0.0050 |
| AZ | 23 | 17 | 15 | 0.925 | 0.0050 |

**Table 3.** mtDNA divergence indices for N. American CPB. Upper half of matrix is average number of nucleotide differences between populations, lower is the net (corrected) nucleotide substitutions per site between populations.

|  | Mex+AZ | Plains | Pest |  |  |  |
| --- | --- | --- | --- | --- | --- | --- |
| Mex+AZ |  | 23.033 | 23.261 |  |  |  |
| Plains | 0.034 |  | 2.734 |  |  |  |
| Pest | 0.035 | 0.00034 |  |  |  |  |
|  | Mex | AZ | Pest | Plains | Mex1 | Mex2 |
| Mex |  | 7.23 |  |  |  |  |
| AZ | 0.00562 |  | 20.088 | 19.904 | 9.535 | 3.930 |
| Pest |  | 0.03225 |  | 2.734 | 26.383 | 20.285 |
| Plains |  | 0.03135 | 0.00034 |  | 26.098 | 20.127 |
| Mex1_GUMROXJL |  | 0.01247 | 0.04378 | 0.04268 |  | 8.652 |
| Mex2_TLTXPB |  | 0.00113 | 0.03260 | 0.03175 | 0.0109 |  |

### AFLP (SP1)

The 130 genotyped individuals yielded a total of 136 polymorphic AFLP bands. The correlation between fragment sizes and their frequencies (-0.09) was not different from zero (P = 0.320). AMOVA showed that the majority of the AFLP variation occurred within sample locations (84.1%) with only 15.9% occurring among populations. Results of AMOVA performed on the three groups; Pest, Plains, and Arizona, were similar, with 92.6 % of the AFLP variation occurring within the groups and 7.4% occurring among groups. ARLEQUIN estimates of the mean within-group AFLP distances (±SD) were similar for the Pest 39.3 (17.32), Plains 38.43 (17.08), and Arizona 34.39 (16.6) groups respectively, with similar mean gene diversity measures as well (Table 4). The sample from Idaho had the lowest diversity (mean within group distance was 28.2 (13.5) and the sample from Washington, which had low mtDNA diversity, had an intermediate level of AFLP diversity, 33.3 (16.6). Using the locus-by-locus analysis in AFLP-surv, the mean genetic diversity (H) using the Lynch and Milligan (1994) method, of the Pest (0.276), Plains (0.272) and Arizona (0.267) samples were very similar as well (Table 4).

**Table 4.** AFLP analysis of genetic diversity within Structure-based populations and Nei’s gene diversity (after Lynch and Milligan 1994).

| Samples | Pairwise Differences ($\theta\pi$): Mean (SD) | Mean Gene Diversity Over Loci (SD) | Theta-S (SD) | Nei's gene Diversity (Expected Heterozygosity $H_j$) (SE). |
| --- | --- | --- | --- | --- |
| Pest | 39.30 (16.58) | 0.29 (0.14) | 27.76 (6.93) | 0.28 (0.01) |
| Plains | 38.43 (17.08) | 0.28 (0.14) | 29.99 (9.06) | 0.27 (0.01) |
| Arizona | 34.39 (16.59) | 0.25 (0.14) | 30.91 (13.16) | 0.27 (0.02) |

Both measures of divergence included in our distance matrix, Nei’s genetic distance calculated using the Lynch and Milligan (1994) method and the mean (corrected) pairwise difference between groups of haplotypes from ARLEQUIN, showed that all Arizona samples were quite distant from the Pest and Plains groups, but that the Pest and Plains groups were similar to each other (Table 5). The STRUCTURE analysis of AFLP haplotypes resulted in three clusters using the Delta K and the Prob(K) estimators. The Pest samples were in one cluster, the Plains samples in a second, and the Arizona samples assigned to a third (Figure 4A). The Pest and Plains clusters showed evidence of some admixture, whereas the Arizona samples showed no admixture with either of the other clusters.

**Table 5.**
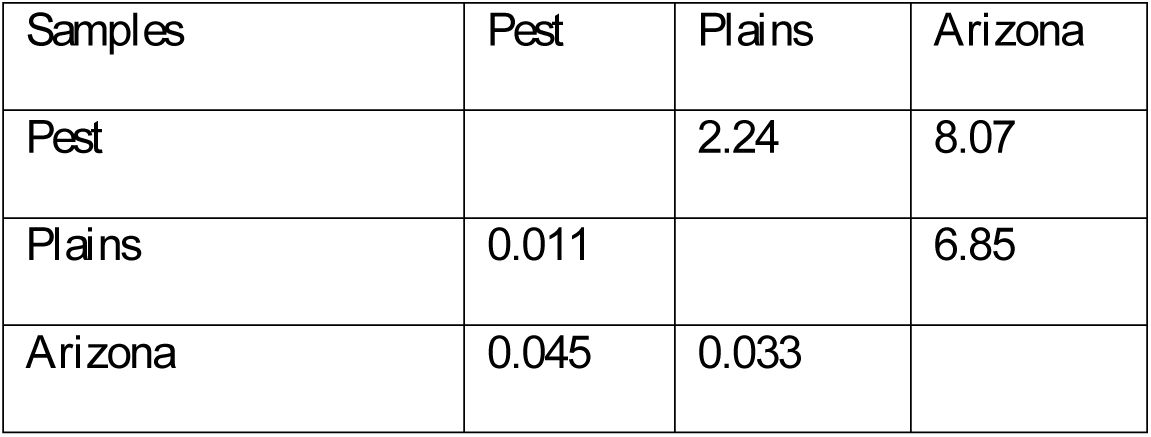
AFLP analyses of genetic distance among U.S. sample regions. Genetic distances among STRUCTURE-based populations presented in two ways: Above diagonal, corrected average number of pairwise distances, below diagonal, Nei’s genetic distances (after Lynch and Milligan 1994)

| Samples | Pest | Plains | Arizona |
| --- | --- | --- | --- |
| Pest |  | 2.24 | 8.07 |
| Plains | 0.011 |  | 6.85 |
| Arizona | 0.045 | 0.033 |  |

**Figure 4.**
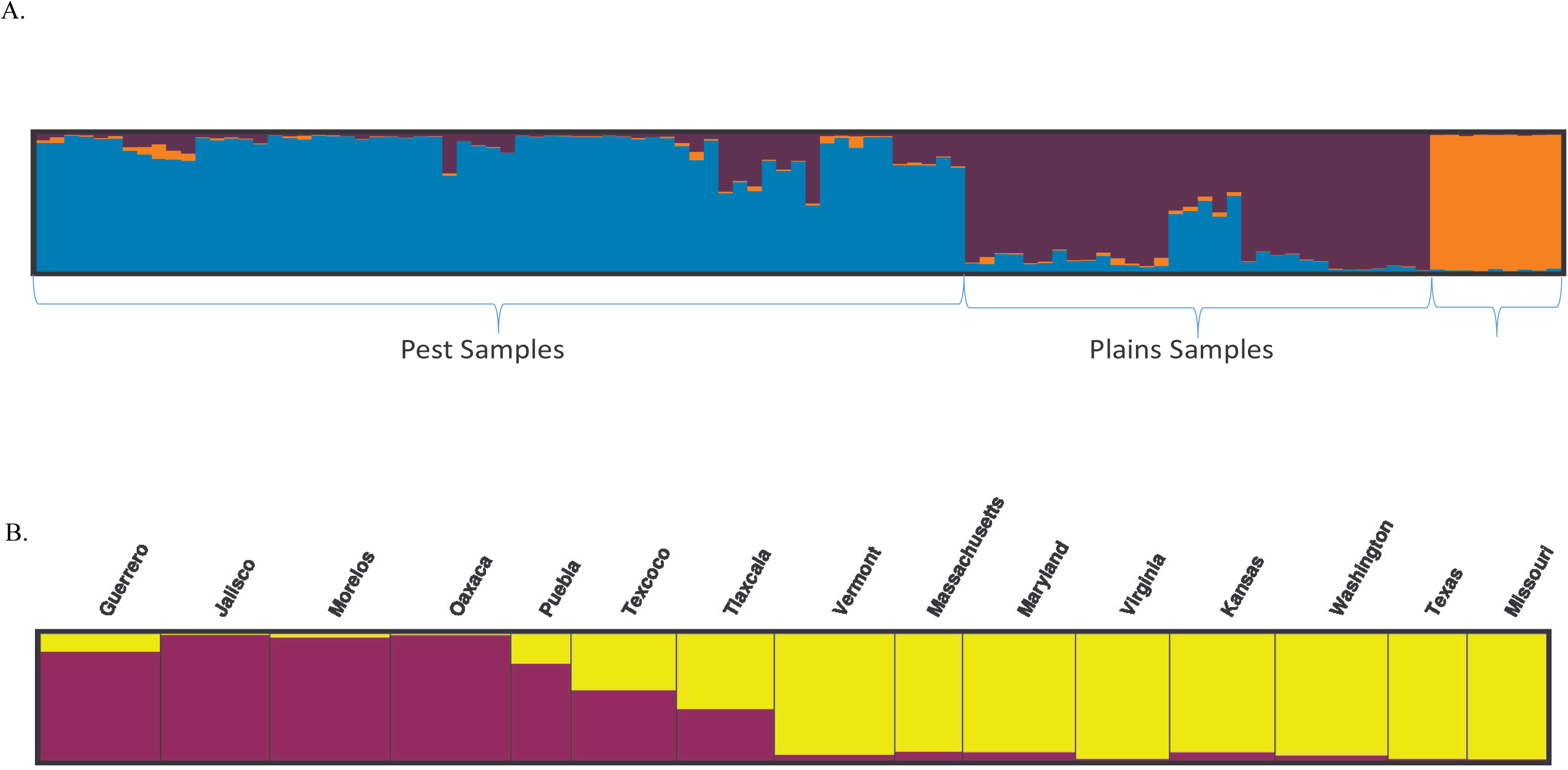
STRUCTURE analysis of AFLP (4A) and microsatellite (4B) data from Colorado potato beetle samples.

### Microsatellites (SP2)

Microsatellite diversity measures among all SP2 samples indicated high diversity across all loci and all sampled populations, RS = 5.25, Theta = 0.162 (0.023), Rst = 0.316 and Fis = 0.009 (0.175). The Neighbor-joining tree clearly partitioned Mexican or U.S. sample locations into two genetically distinct groupings (Figure 5), with Mexican populations divided into the Groups 1 and 2 observed by the mtDNA analysis. Pairwise comparisons of Nei’s genetic distances among sample locations uncovered large divergence among many populations, but little difference between some neighboring population pairs such as Texcoco-Tlaxcala and Virginia-Maryland (Figure 5). Microsatellite based genetic distances between the two clades of Mexican samples and Pest and Plains samples reveal large genetic distances between Mexican and Pest + Plains samples and much smaller distances between Pest and Plains samples (Table 6). Microsatellite diversity within these regional samples was nominally largest within the Plains sample and less in the individual Mexican samples, though differences among the regions were not significant (Table 7).

**Table 6.** Genetic distances among geographic groupings. Microsatellite analysis of genetic distances among regional populations. Nei’s Chord distance as calculated in FSTAT (Goudet 1995).

| Sample | Mexico 2 | Pest | Plains |
| --- | --- | --- | --- |
| Mexico 1 (GUMROXJA) | 0.118 | 0.289 | 0.347 |
| Mexico 2 (TLTXPB) |  | 0.213 | 0.272 |
| Pest |  |  | 0.085 |

**Table 7.** Microsatellite Analysis of genetic diversity within sample groupings from Mexico and Pest and Plains samples in the U.S. Four microsatellite loci were usable for the MEX1 sample and three were usable for the remainder of the populations.

| Sample | N | Mean Gene Diversity | Theta-H | Nei's gene Diversity (Expected Heterozygosity $H_j$) (SE). |
| --- | --- | --- | --- | --- |
| Mexico 1 (GUMROXJA) | 224 | 0.692(0.407) | 1.673 | 0.641 (0.251) |
| Mexico 2 (TLTXPB) | 166 | 0.584 (0.377) | 1.903 | 0.705 (0.144) |
| Pest | 304 | 0.716 (0.447) | 1.972 | 0.719 (0.124) |
| Plains | 106 | 0.725 (0.449) | 2.126 | 0.744 (0.122) |

**Figure 5.**
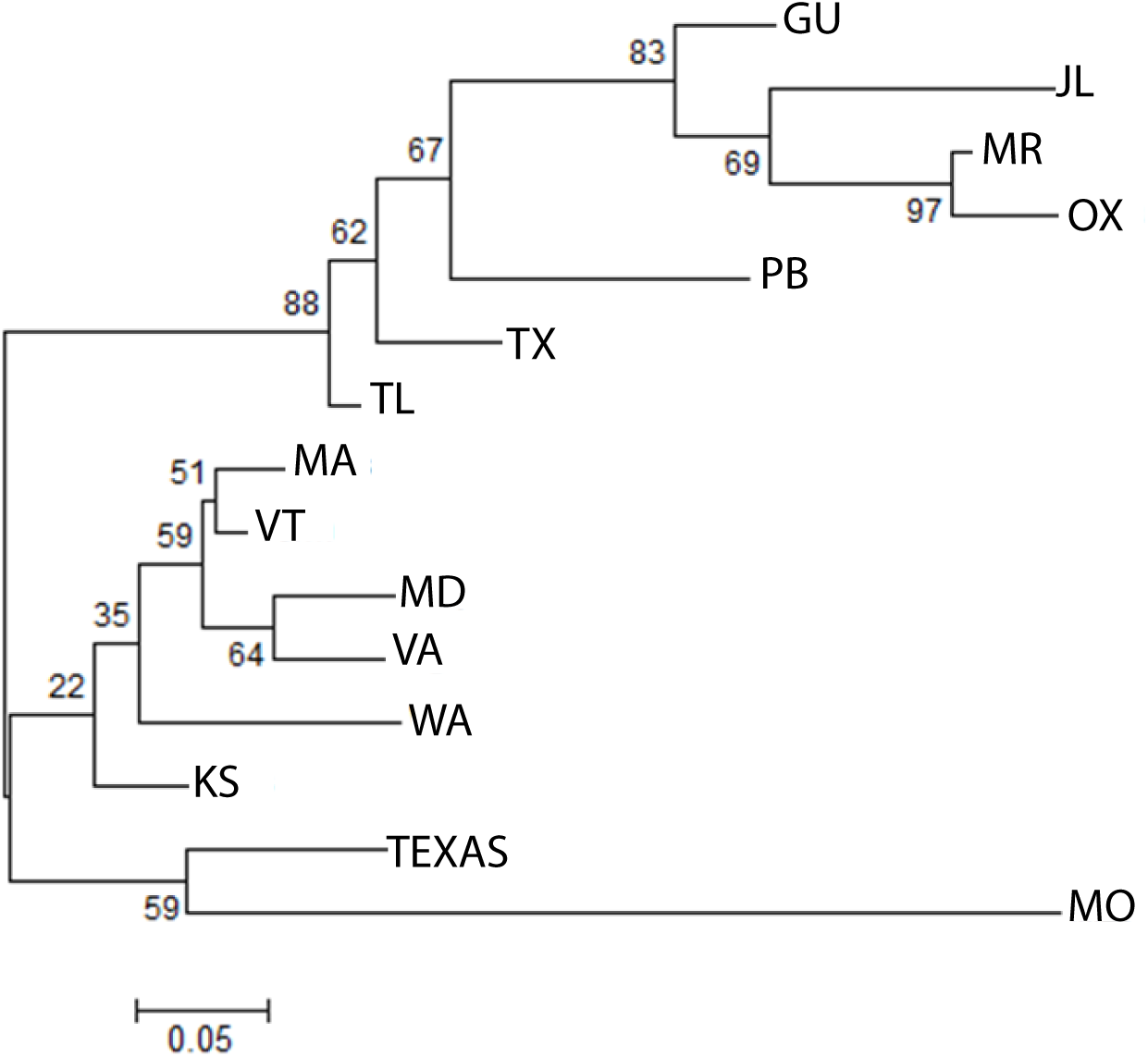
Neighbor-joining phylogram of microsatellite genotypes from Colorado beetle. Nodes are labelled with bootstrap support from 10,000 replicate phylogram constructions.

Similar to our distance-based analyses of the microsatellite data (Table 6, Figure 5), the STRUCTURE analysis resulted in two clusters, with U.S. Pest and Plains samples in one cluster and Mexico samples in the other (Figure 4B). STRUCTURE results for K = 2 resulted in some beetles from the Central Highlands of Mexico (Texcoco, and Tlaxcala) having membership in both groups. Similarly, some U. S. Pest and Plains individuals were placed in the Mexico cluster. Because a limited number of loci was used in this analysis, it is difficult to distinguish between shared recent ancestry of some individuals within populations and mis-categorization of individuals by STRUCTURE.

## Discussion

Despite the widespread distribution of *L. decemlineata* throughout the Northern Hemisphere, the geographic and evolutionary origins of pest populations of the beetle have long been obscure. We used three types of molecular marker to test whether contemporary genetic patterns support one of three proposed hypotheses: the Out-of-Mexico hypothesis, the Hybrid Origin hypothesis, or the Plains Native hypothesis. Overall, the population genetic and phylogeographic analysis provide the greatest support for the Plains Native hypothesis. All three sets of genetic markers used in this study show a large phylogenetic divide between the Plains + Pest beetles and the Mexico + Arizona beetles. None of the mtDNA haplotypes were shared by Mexico + Arizona and Plains + Pest samples. Furthermore, the two clades showed a high level of divergence. The closest Plains haplotypes were at least 15 nucleotide substitutions (out of 577 nucleotides per haplotype) different from the Mexican or Arizona haplotypes. We calculated that the mean COI-COII nucleotide divergence between Plains and Pest samples and the Mexico + Arizona samples (corrected for intra-population divergence) was 3.5%, representing ca. 1 million years of divergence (Papadopoulou et al. 2010). In addition, CPB populations from the southern plains of the U.S. have similar mitochondrial haplotype diversity as samples from Arizona and of the two Mexican clades of beetles, indicating that Plains beetles are not recent colonists. The Plains samples also had slightly greater mitochondrial haplotype diversity than the Pest samples, as expected if the pest populations originated from a subsample of those in the Plains. Both sets of nuclear markers (AFLP and microsatellites) also showed a significant genetic divergence between the Mexico + Arizona and the Plains + Pest samples (Figures 4 & 5, Tables 6 & 7) and neither demonstrated the large decrease in genetic diversity in the Pest samples that would be expected following a recent introduction from central Mexico (Table 1). Based on the mitochondrial data, we suggest that there has been limited to no gene flow between the Mexico + Arizona and the Plains + Pest clades and there is no evidence of a colonization-mediated genetic bottleneck.

These genetic data support the idea that Pest beetles are derived from Plains beetles, in agreement with the oldest hypothesis for their origin and the historical documentation of the beetle’s expansion, from the eastern slope of the Rocky Mountains across the U.S. following its shift onto potato (Walsh 1865, Riley 1877, Tower 1906, Casagrande 1985, Hsiao 1985). Beetles from the Washington and Idaho samples are genetically similar to other beetles in the Plains + Pest group, though possibly less variable, suggesting that the founders were long-distance colonists from Great Lakes or eastern U.S. populations as suggested by contemporaneous records (Tower, 1906).

The complete mismatch in mitochondrial DNA haplotypes between the Mexican + Arizona and Plains + Pest subgroups clearly does not support the Out-of-Mexico hypothesis. Similarly, the neighbor-joining tree of the microsatellites and the distance matrix of the AFLPs show similarity of Pest and Plains samples and large phylogenetic gaps between Mexican + Arizona and Plains + Pest subgroups, casting further doubt in the Out-of-Mexico hypothesis. However, the STRUCTURE analysis of the microsatellite data, though limited here because of few loci and the high likelihood of homoplasy, shows some genetic similarity of the TX-TL-PB group of Mexican samples, as also observed with the mitochondrial fragment analyses (Figures 2 & 3). Interestingly, in this analysis, some of those samples cluster to a small degree with some U. S. samples. Those three states are in the Central Highlands of Mexico, where beetles were collected off of *S. rostratum* and the sites were at higher altitudes than the other collections. Many fewer of the individuals with membership in both clusters were collected off of *S. angustifolium* in the neighboring states of Morelos and Guerrero, which are also lower in altitude, suggesting two possible environmental differences (altitude and host plant) that may underlie the divergence among Mexican beetle populations. The predominant conclusion from all markers and analyses presented here is that we can reject the Out-of-Mexico hypothesis for the origin of Pest populations of CPB.

Our data do not support the Hybrid Origin hypothesis (Table 1), where descendent Pest populations should show greater levels of genetic diversity than Plains and Mexican beetles. Pest populations did not have greater diversity, using any of the three genetic marker types, which would be expected if they are the product of divergent populations that had hybridized. Similarly, as Pest populations were much more similar to the Plains populations than to any other group, they were not genetically intermediate to the Plains and Mexico + Arizona populations as expected from a hybridization scenario. Pest populations were also not genetically divergent from the Plains samples as would occur following hybridization of plains individuals with an unsampled “ghost” population or species.

It appears that the process of colonization of potato as a host plant and agroecosystem may have occurred with extensive gene flow from populations on *S. rostratum* in the Central Plains of the U.S. onto potato as we found seven mitochondrial haplotypes that occur in both Pest and Plains samples. Using the haplotype network and the criteria for determining founder haplotypes described in Richards *et al.* (2000), potential colonist haplotypes should be shared among Pest and Plains samples, and have immediate derivative haplotypes that are found only in Plains samples (e.g., haplotypes 1, 5, 3), to distinguish those that are shared possibly because of recent back-migration. Because the founder analysis is dependent on the scale and resolution of the sampling and on the accuracy of the haplotype network, we resist putting too much confidence in the inference of any single founder. These data suggest, as noted above, that multiple founder haplotypes can still be found in the ancestral (pre-potato) geographic ranges in the southern plains of the U.S. Additional sampling, especially in the highly variable plains region, would likely increase the number of candidate founder haplotypes.

If pest populations of CPB originated from resident populations in the plains region, then additional studies are needed to resolve the ancestral host plant of those populations. It is unclear that the beetle’s ancestral host plant, *S. rostratum* is native to the plains region. Plains populations of the beetle’s host plant, *S. rostratum* show signs of a recent population bottleneck compared to populations in Mexico, suggesting that *S. rostratum* is a recent colonist of the plains (Zhao et al. 2013). However, a STRUCTURE analysis of *S. rostratum* microsatellites also predicted two populations, in U. S. plains states and the Central Highlands of Mexico (Zhao et al. 2013), which could indicate that the plant has a long residency in the Plains states. Because the beetle feeds on an array of wild plants within Solanaceae (Hare and Kennedy 1986, Jacques 1988, Hare 1990), it is possible that Plains beetles were associated with other host plants as well as *S. rostratum* before the arrival of potato.

Colorado potato beetle has displayed an extraordinary ability to adapt to a wide range of host plants, climates, and insecticides (Hare 1990, Alyokhin et al. 2008b, Lehmann, et al. 2015, Hiiesaar et al. 2016), which has enabled it to colonize practically all potato-growing regions in the Northern Hemisphere (Weber 2003, Liu et al. 2012). Our data provide clear evidence that CPB expanded onto potato from *S. rostratum* on the Central Plains of the U.S. The host expansion appears to have occurred without a genetic bottleneck, suggesting considerable gene flow between beetles on potato and *S. rostratum*, at least early in the invasion process. This improved understanding of the ancestry of pest populations will guide sampling of CPB to determine the origins of genetic variation in traits relevant to the beetle’s success. We are also better able to assess whether those genetic changes were due to novel mutation or standing genetic variation in large genetically diverse populations. By understanding the source of contemporary pest populations and some of the dynamics associated with the colonization of potato, we improve our ability to analyze the genetic, ecological and evolutionary dynamics associated with CPB adaptation to the potato agroecosystem.

## Acknowledgements

Andrea Badgley, Darren Schnieder, Joe Labrum assisted with laboratory work, Todd Peters, Damon Orsetti, Juan Nuñez-Farfan, Segura Obdulia-Leon, P Michaud, Carolina Lukac and Celso Morales helped with collections, and Steve Lingenfelter, Sonya Scheffer and Charlie Mitter contributed through discussions of early drafts of this manuscript.

